# Targeting the host transcription factor HSF1 prevents human cytomegalovirus replication in vitro and in vivo

**DOI:** 10.1101/2024.09.23.614483

**Authors:** Dilruba Akter, Juthi Biswas, Michael J. Miller, Dennis J. Thiele, Eain A. Murphy, Christine M. O’Connor, Jennifer F. Moffat, Gary C. Chan

## Abstract

FDA-approved antivirals against HCMV have several limitations, including only targeting the later stages of the viral replication cycle, adverse side effects, and the emergence of drug-resistant strains. Antivirals targeting host factors specifically activated within infected cells and necessary for viral replication could address the current drawbacks of anti-HCMV standard-of-care drugs. In this study, we found HCMV infection stimulated the activation of the stress response transcription factor heat shock transcription factor 1 (HSF1). HCMV entry into fibroblasts rapidly increased HSF1 activity and subsequent relocalization from the cytoplasm to the nucleus, which was maintained throughout viral replication and in contrast to the transient burst of activity induced by canonical heat shock. Prophylactic pharmacological inhibition or genetic depletion of HSF1 prior to HCMV infection attenuated the expression of all classes of viral genes, including immediate early (IE) genes, and virus production, suggesting HSF1 promotes the earliest stages of the viral replication cycle. Therapeutic treatment with SISU-102, an HSF1 inhibitor tool compound, after IE expression also reduced the levels of L proteins and progeny production, suggesting HSF1 regulates multiple steps along the HCMV replication cycle. Leveraging a newly developed human skin xenograft transplant murine model, we found prophylactic treatment with SISU-102 significantly attenuated viral replication in transplanted human skin xenografts as well as viral dissemination to distal sites. These data demonstrate HCMV infection rapidly activates and relocalizes HSF1 to the nucleus to promote viral replication, which can be exploited as a host-directed antiviral strategy.

**One Sentence Summary:** Inhibiting of HSF1 as a host-directed antiviral therapy attenuates HCMV replication in vitro and in vivo.

## INTRODUCTION

Human cytomegalovirus (HCMV) is an endemic β-herpesvirus with a prevalence of 60 - 90% among adults depending on geographical and socioeconomic status (*1, 2*). HCMV infection can lead to a wide range of clinical manifestations. Although generally asymptomatic in healthy individuals, HCMV is a causative agent of a significant percentage of infectious mononucleosis (*3*). HCMV is also linked to several chronic inflammatory diseases, such as atherosclerosis and inflammatory bowel disease (*4, 5*), and certain cancers, such as glioblastoma and breast cancer (*6, 7*). Patients with compromised immunity, including organ recipients, chemotherapy patients, AIDS patients, or neonates, are at risk of experiencing severe and potentially life-threatening complications from an acute HCMV infection (*8*). In these patients, the broad tissue and cellular tropism of HCMV can lead to a wide range of organ pathologies and ultimately end-organ disease (*9*).

Upon HCMV entry into susceptible and permissive cell types, three temporal classes of viral lytic genes are expressed in a tightly controlled manner: immediate early (IE), early (E), and late (L) genes (*10, 11*). After replication of viral DNA, the synthesis of structural proteins triggers the assembly of the capsid followed by egress of progeny viruses (*11*). Currently licensed antiviral drugs ganciclovir, valganciclovir, foscarnet, cidofovir, letermovir, and maribavir all target viral factors that facilitate critical post IE steps of the lytic replication cycle (*12, 13*), which is likely due to screening assays relying on the measuring of viral genome replication. However, despite selectively targeting viral proteins, the prolonged use of these antivirals is associated with severe drug toxicities (*14*). As such, treatment duration is limited with rebound infection and subsequent disease often occurring upon termination of the drug regimen in high-risk patients (*15, 16*). Further complicating the treatment of HCMV infections is the emergence of drug resistant HCMV strains (*12, 17*). Thus, there is an existing need for the development of anti-HCMV drugs that target different stages of the lytic replication cycle, exhibit reduced side effects, and have an increased barrier to the development of drug-resistant mutations.

The targeting of host factors rather than viral proteins has emerged as an alternative antiviral strategy that offers the benefit of an increased threshold to viral resistance due to their role in mediating normal cellular functions (*18, 19*). Unfortunately, host-directed antiviral therapeutics is also troubled with the possibility of unwanted side effects and/or cytotoxicity (*18*). A potential approach to circumvent the toxicity of host-directed drugs is the targeting of cellular factors required for viral infection that are activated specifically in infected cells. The heat shock (HS) response is a cellular protective mechanism activated in response to different stresses. Importantly, HCMV has evolved to usurp many HS-responsive proteins, including the heat shock proteins (HSPs) HSP40, HSP70, and HSP90, to directly promote lytic replication, virion protein folding and assembly, and host cell survival (*20–23*). HSPs are regulated by the stress responsive transcription factor heat shock transcription factor 1 (HSF1), which remains inactive in a monomeric form complexed with HSPs in the cytoplasm under homeostatic conditions. During times of stress, HSPs release HSF1 allowing for trimerization of HSF1 followed by translocated into the nucleus to promote DNA binding and transcription from heat shock element (HSE) containing promoter regions (*24*). Recent findings have demonstrated HSF1 activity promotes the replication of human immunodeficiency virus (HIV), Dengue virus, coxsackievirus B3, vaccinia virus (VACV), Epstein–Barr virus (EBV), and human coronaviruses (*25–29*). To date, whether HSF1 has any role in promoting HCMV replication remains unknown.

Our study revealed that HCMV infection triggered a non-canonical, bi-phasic activation of HSF1 when compared to the canonical transient activation induced by HS. To investigate the role of HCMV-induced HSF1 activation, we utilized SISU-102, a tool compound validated to directly bind and selectively inhibit nuclear HSF1 activity. A previous study also showed that SISU-102 was efficacious, inhibited HSF1 activity, and was well tolerated in murine cancer models (*30*). We found SISU-102, or genetic inhibition of HSF1, attenuated HCMV IE and L gene expression and ultimately progeny production while having minimal effect on host cell viability. Further, SISU-102 was effective against multiple HCMV strains, exhibiting a similar in vitro potency as the standard-of-care drug ganciclovir. Because of the strict species specificity of HCMV, in vivo models are limited to expensive humanized murine models investigating latency or murine models with non-biologically relevant settings to examine active lytic replication. To circumvent these limitations and test the therapeutic potential of inhibiting HSF1, we utilized a human skin xenograft transplant murine model originally developed by our group to investigate acute HCMV infection in biologically relevant tissue in vivo. We found that SISU-102 was more effective at attenuating HCMV replication at initial sites of infection as well as preventing viral dissemination to distal uninfected sites when compared to a standard-of-care drug valganciclovir, with no observable adverse side effects. Thus, this study provides a proof-of-concept for HSF1 as a new druggable cellular target to block HCMV replication and dissemination.

## RESULTS

### HCMV stimulates a bi-phasic, persistent activation of HSF1 during lytic infection

A central player regulating the HS response is the stress response transcription factor HSF1 (*31–35*). Although HCMV hijacks several HSPs to promote lytic replication (*20–23*), the role of HSF1 in facilitating HCMV infection remains to be determined. Thus, we examined if HSF1 is activated during a lytic infection. Fibroblasts were infected with HCMV strain TB40E over a time course of infection and HSF1 phosphorylation levels measured at serine 326 (Ser^326^), a hallmark of activation (*36*) (**Fig. 1A, 1B**). Phosphorylated HSF1 was observed as early as 30 minutes post infection (mpi) and maintained through 4 hours post infection (hpi) until a secondary burst at 24 hpi occurred. In contrast, fibroblasts infected with UV inactivated-HCMV (UV-HCMV) stimulated an early (30 mpi) but not late (24 hpi) phosphorylation of HSF1, suggesting the initial activation of HSF1 is due to the viral entry process and the secondary activation is mediated by de novo synthesized viral gene products. Similar to fibroblasts infected with UV-HCMV, HS-treated cells exhibited an early transient activation of HSF1 at 30 minutes (min) post treatment that dissipated by 24 hours (h). Further, HS also increased the apparent molecular weight of HSF1 relative to HCMV infection, indicating the presence of distinct post-translational modifications on HSF1 when activated by HCMV infection versus HS. The activation of HSF1 by HCMV strain TB40E was not strain specific as HCMV strain Towne also stimulated a rapid and persistent activation (**Fig. 1C, 1D**). Thus, HCMV induces a bi-phasic activation of HSF1, suggesting HCMV is actively targeting and modifying HSF1 activity rather than HSF1 activation simply being a cellular response to a viral infection.

**Figure 1.**
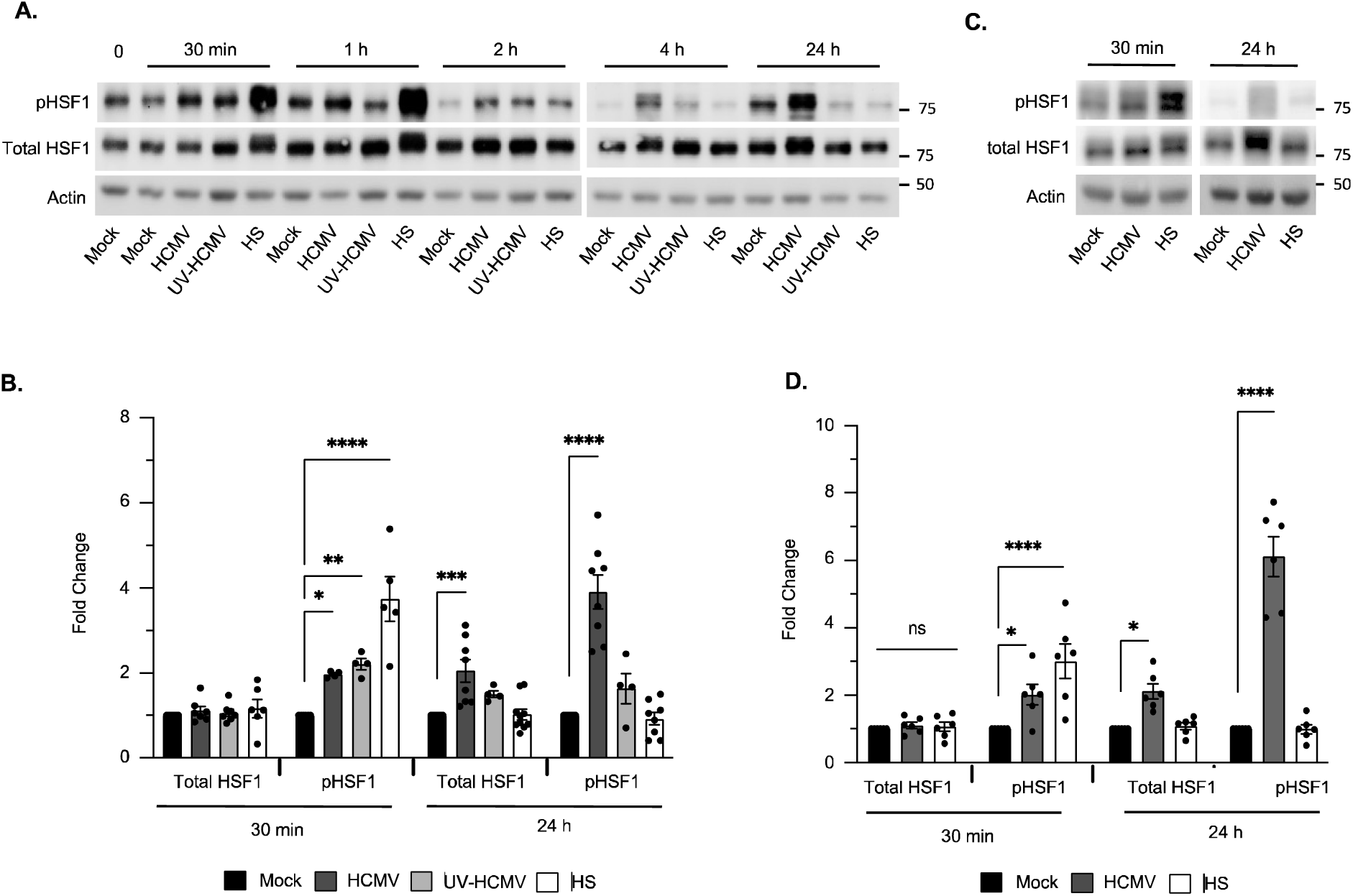
HCMV induces a persistent activation of HSF1 during lytic infection. **(A to D)** HEL 299 fibroblasts were mock infected or infected (MOI 5) with HCMV strain TB40E (A, B), UV-inactivated HCMV strain TB40E (UV-HCMV) (A, B), or HCMV strain Towne (C, D) for indicated time points. As a positive control, fibroblasts were subjected to heat shock (HS) for 15 min at 42°C. Total HSF1 and phosphorylation of HSF1 at Ser^326^ were detected by Western blot (A, C) and quantified (B, D). β-actin was used as a loading control. Western blots and densitometry are representative of at least 3 biological replicates per group. ns, non-significant, *P<0.05, **P□<□0.005, ****P□<□0.0001, by one-way ANOVA with Tukey’s HSD post hoc test.

### Prophylactic inhibition of HSF1 attenuates HCMV IE protein abundance

Next, we sought to determine whether the noncanonical activation of HSF1 during early HCMV infection promotes viral gene expression. Fibroblasts were treated for 24 h with increasing concentrations of SISU-102, which directly binds to the amino-terminal HSF1 DNA binding domain to stimulate degradation of active nuclear HSF1 through a well-established ubiquitin-proteasome-dependent pathway (*30, 37*). Fibroblasts treated with SISU-102 were then infected for 24 h with IE2-eGFP, a recombinant HCMV stain TB40E engineered to express mCherry under a constitutively active SV40 promoter and eGFP-tagged viral immediate early 2 (IE2) protein under the control of the viral major immediate early promoter (MIEP). Because IE2 function is attenuated by fusion with eGFP, a self-cleavage T2A linker peptide was placed in-between the two proteins to allow for IE2 activity (*38*). The use of IE2-eGFP allows for the rapid evaluation of the consequences of SISU-102 administration prior to genome replication, and thus represents an important tool for the future screening of novel antivirals that act at the immediate early stages of infection. Following infection of fibroblasts with IE2-eGFP, HSF1 accumulated in the nucleus and, as expected, SISU-102 treatment strongly reduced the accumulation of nuclear HSF1 (**fig. S1A**), while having no effect on HCMV binding and entry into fibroblasts (**fig. S2A, S2B**). At 24 hpi, we found a dose-response reduction in IE2 abundance where the half-maximal inhibitory concentration (IC_50_) of SISU-102 was ∼ 8.211 µM (**Fig. 2A**). SISU-102 exhibited minimal cytotoxicity at 8.211 µM (**Fig. 2B**), suggesting the reduction of IE2 was not due to reduced cell viability. As expected, ganciclovir, had no effect on IE2 levels or cell viability as it prevents the incorporation of nucleotides into elongating viral DNA to block post IE replication steps (**Fig. 2C, 2D**) (*39, 40*). Western blot analysis also showed SISU-102 reduced the levels of the viral IE1 protein, indicating that SISU-102 has broad impact on IE protein abundance (**Fig. 2E**). To further validate our pharmacological studies, we depleted HSF1 abundance utilizing an HSF1-specific siRNA, which resulted in ∼90% knockdown (**Fig. 2F**). Consistent with SISU-102, HSF1-depleted infected fibroblasts had reduced IE1 levels at 24 hpi. Knockdown of HSF1 did not stimulate cell death (**Fig. 2G**), confirming that the reduction of IE proteins due to HSF1 inhibition was not an indirect effect on cell viability. These data suggest HSF1 could serve as an antiviral cellular target to suppress HCMV lytic replication.

**Figure 2.**
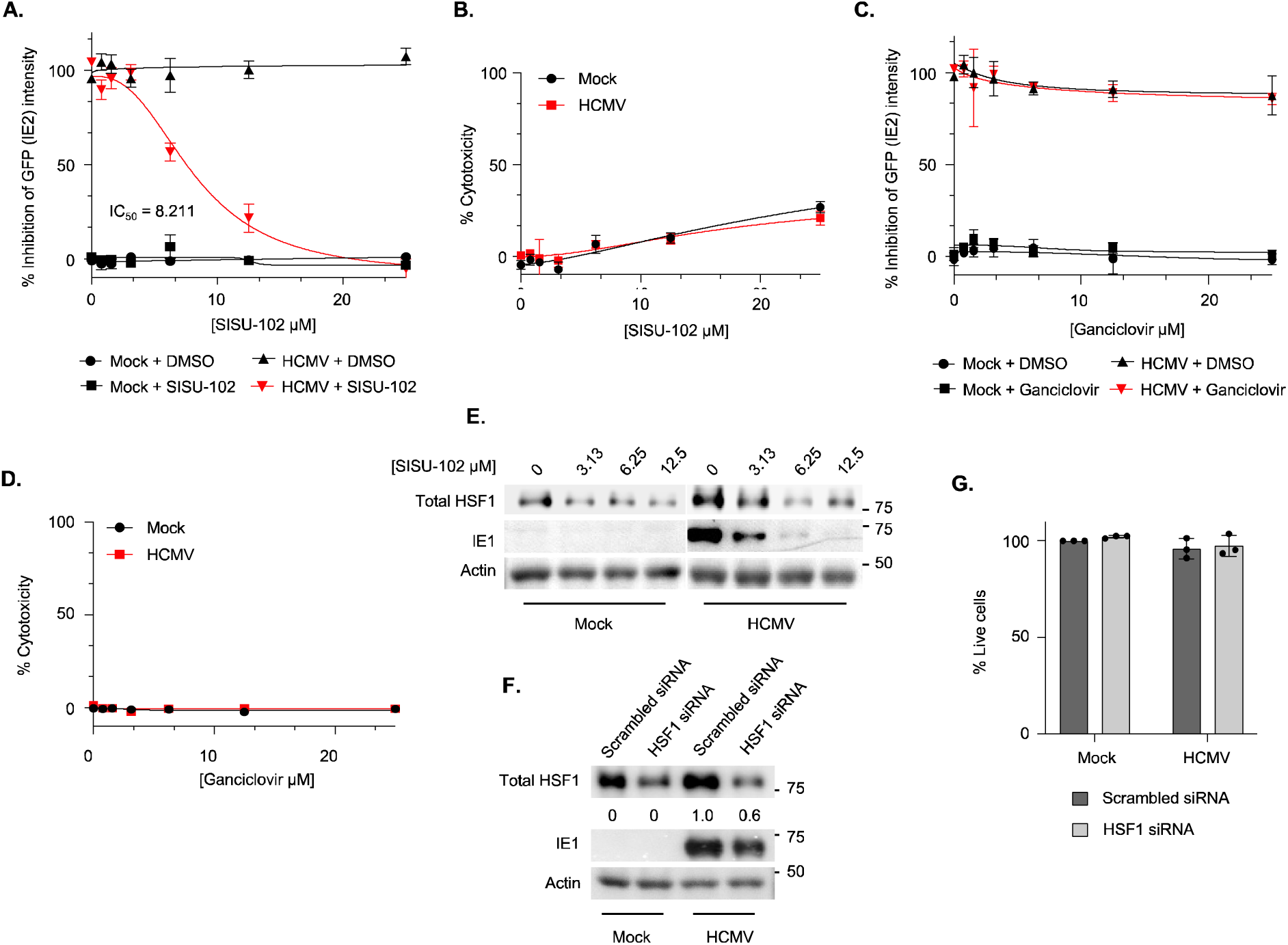
Inhibition of HSF1 reduces HCMV IE protein abundance. **(A to E)** HEL 299 fibroblasts were prophylactically treated with SISU-102 (A, B, E) or ganciclovir (C, D) at the indicated concentrations for 24 h. **(F, G)** Fibroblasts were transfected with a scrambled or HSF1 siRNA for 48 h. Following drug treatment or siRNA knockdown, cells were mock infected or infected (MOI 1) with IE2-eGFP for an additional 24 h. GFP intensity was measured using a fluorescent plate reader (A, C). Cytotoxicity was determined by SRB colorimetric assay (B, D, G). Total HSF1 and IE1 were detected by Western blot (E, F). IE1 was quantified by densitometry (F). β-actin was used as a loading control. All data are representative of at least 3 biological replicates per group.

### Prophylactic inhibition of HSF1 reduces HCMV L proteins and virus progeny production

IE proteins are required for the subsequent expression of viral E and L genes (*11*). To test whether that the decrease in IE proteins mediated by SISU-102 causes the expected reduction of L proteins, we engineered a recombinant virus that expresses mCherry from a constitutively active SV40 promoter and the viral L protein UL99 fused to eGFP driven by a true late promoter (UL99-eGFP) (*41*). Fibroblasts were prophylactically treated with SISU-102 for 24 h, followed by infection with UL99-GFP for 96 h. As with IE protein abundance, UL99 levels exhibited a dose-dependent reduction in response to increasing concentrations of SISU-102 with an IC_50_ value of ∼3.898 μM (**Fig. 3A**). SISU-102 had no impact on cell viability at ∼3.898 μM (**Fig. 3B**), suggesting the negative impact of SISU-102 on L protein levels was due to HSF1 inhibition, rather than on the viability of infected cells. SISU-102 had a similar efficacy as ganciclovir (IC_50_ ∼2.683 μM) (**Fig. 3C**), which also had no effect on cell viability (**Fig. 3D**). However, the potency of ganciclovir leveled off rapidly at concentrations above its IC_50_, never completely abolishing UL99 levels. In contrast, SISU-102 continued to reduce UL99 abundance to near baseline levels. A possible explanation for SISU-102’s enhanced potency relative of ganciclovir at concentrations beyond its IC_50_ could be due to the inhibition of multiple viral kinetic gene classes. To further control for the potential off-target effects of SISU-102, siRNA-mediated knockdown of HSF1 was performed and levels of glycoprotein L (gL), a viral L protein, evaluated (**Fig 3E**). HSF1-depleted fibroblasts infected with WT HCMV exhibited decreased gL abundance when compared to control fibroblasts transfected with scrambled siRNA, indicating a global reduction of viral L proteins in the absence of HSF1 activity.

**Figure 3.**
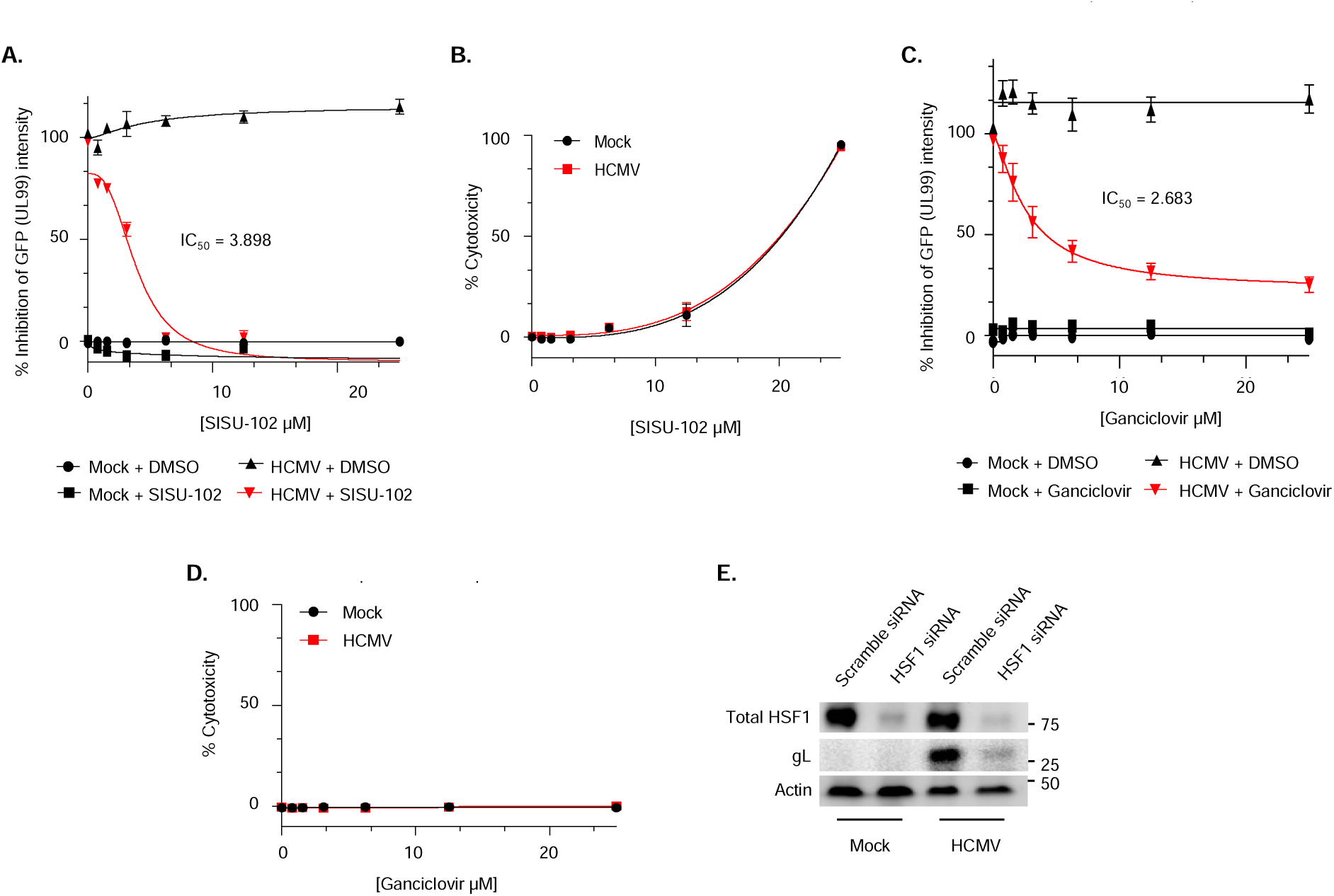
HSF1 inhibition attenuates the levels of HCMV L proteins during lytic replication. **(A to D)** HEL 299 fibroblasts were prophylactically treated with SISU-102 (A, B) or ganciclovir (C, D) at the indicated concentrations for 24 h. **(E)** Fibroblasts were transfected with a scrambled or HSF1 siRNA for 48 h. Following drug treatment or siRNA knockdown, cells were mock infected or infected with UL99-GFP for 96 h at an MOI of 1 (A to D) or 0.1 (E). GFP intensity was measured using a fluorescent plate reader (A, C). Cytotoxicity was determined by SRB colorimetric assay (B, D). Total HSF1 and gL were detected by Western blot (E). β-actin was used as a loading control. All data are representative of at least 3 biological replicates per group.

Next, to test if the reduction of IE and L proteins corresponds to a decrease in viral genome replication and progeny production, we treated fibroblasts with SISU-102 or ganciclovir for 24 h with increasing concentrations starting at 3.13 μM, which represents the approximate IC_50_ values of each inhibitor based on viral L protein abundance (**Fig. 3A, 3C**). After pretreatment with inhibitors, fibroblasts were infected with HCMV strain TB40E for an additional 96 h. qRT-PCR and TCID50 assays were used to measure the amount of cell-associated viral genomes and released progeny virus, respectively. HSF1 inhibition with SISU-102 reduced the levels of cell-associated viral genomic DNA and resulted in a 4-log reduction in progeny production (**Fig. 4A, 4B**). The magnitude of reduction in viral genomes and progeny was similar to ganciclovir treatment. The effects of SISU-102 on HCMV genome replication and virus production were also observed with the HCMV strain Towne (**Fig. 4C**), demonstrating antiviral efficacy of SISU-102 across different HCMV strains. As further confirmation, siRNA knockdown of HSF1 attenuated progeny production (**Fig. 4D**). Thus, HSF1 is necessary for the efficient progression of HCMV’s lytic replication cycle and that prophylactic inhibition of HSF1 significantly attenuates viral replication and the production of progeny virus.

**Figure 4.**
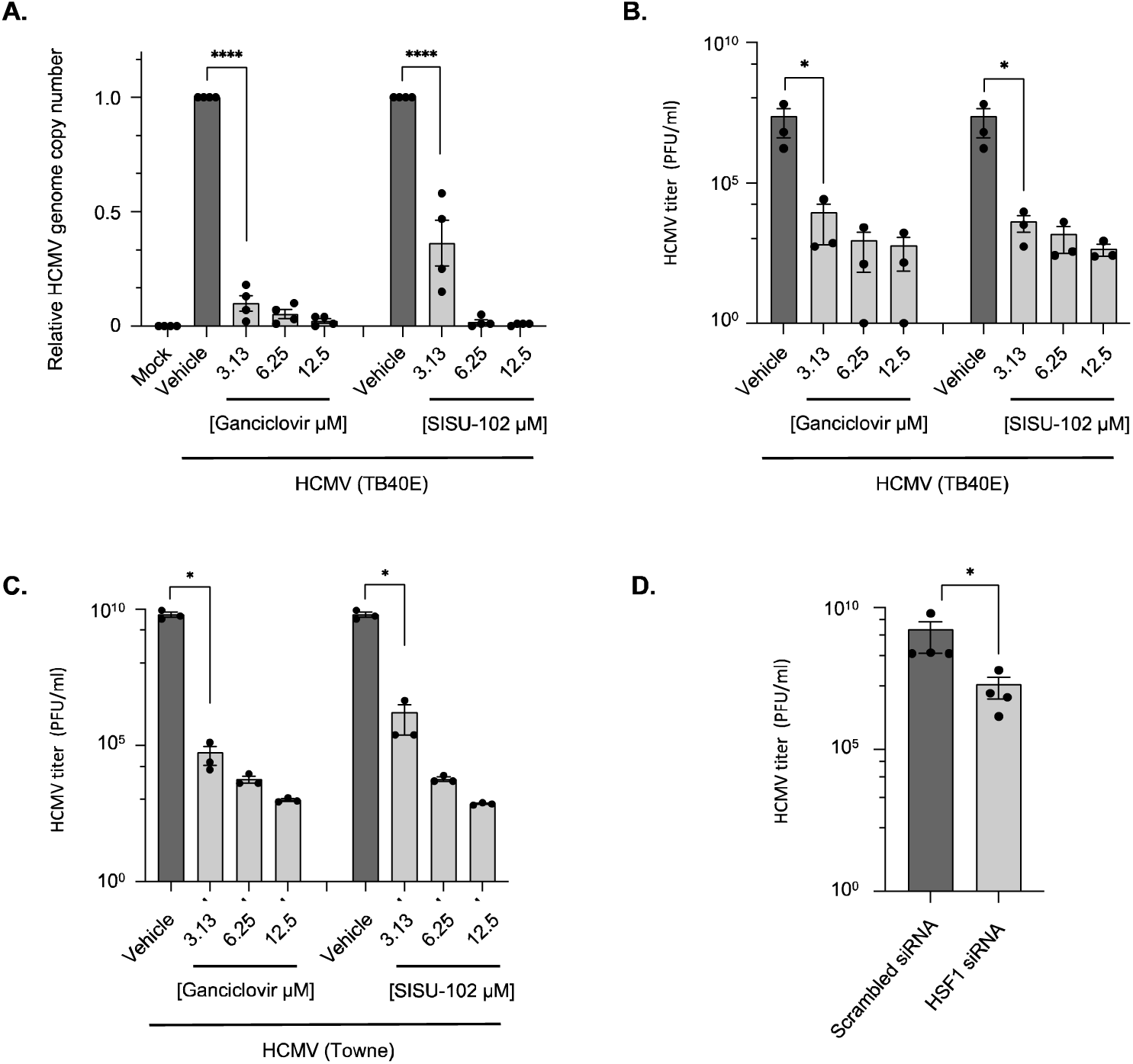
HSF1 activity is required for efficient viral replication and progeny production. **(A to C)** HEL 299 fibroblasts were prophylactically treated with SISU-102 or ganciclovir at the indicated concentrations for 24 h. **(D)** Fibroblasts were transfected with a scrambled or HSF1 siRNA for 48 h. Following drug treatment or siRNA knockdown, cells were mock infected or infected (MOI 1) with WT HCMV strain TB40E (A, B, D) or HCMV strain Towne (C) for 96 h. Viral genome copy number was assessed by qPCR analysis for *UL123* (as a marker of the viral genome) and *GAPDH* (A). TCID_50_ assays were performed on the supernatants to measure progeny virus production (B to D). All data are representative of 3 to 4 biological replicates per group. ****P < 0.0001, by one-way ANOVA with Tukey’s HSD post hoc test (A). *P<0.05, by Mann Whitney t-test (One-tailed) (B to D).

### Therapeutic inhibition of HSF1 reduces HCMV replication and progeny production

Prophylactic or pre-emptive HCMV antivirals are given as standard-of-care to transplant recipients at high-risk of HCMV exposure. However, demonstrating therapeutic effectiveness of targeting HSF1 could impact the prognosis of transplant patients exhibiting a rebound infection once standard-of-care therapies are discontinued or other vulnerable populations such as congenitally infected newborns. Thus, we assessed the therapeutic efficacy of SISU-102 to limit HCMV infection after the lytic replication cycle has already been initiated. To this end, fibroblasts were infected with UL99-GFP for 24 h, after maximal viral IE protein production has been reached, followed by treatment with increasing concentrations of SISU-102 and infection allowed to progress for an additional 72 h. UL99 expression showed a dose-dependent reduction in response to SISU-102 with no effect on the viability of uninfected cells at the IC_50_ of 3.778 μM (**Fig. 5A, 5B**). Surprisingly, HSF1 inhibition reduced the viability specifically of HCMV-infected fibroblasts (**Fig. 5B**), indicating that the viability of pre-infected cells is sensitive to the loss of HSF1 activity and that SISU-102 could selectively eliminate infected cells as well as terminate viral replication. Ganciclovir had a similar therapeutic IC_50_ with no effect on the cell viability of uninfected and infected fibroblasts (**Fig. 5C, 5D**). However, similar to prophylactic treatment (**Fig. 3C**), the potency of therapeutically administered ganciclovir leveled off at concentrations above its IC_50_ whereas SISU-102 reduced UL99 to baseline levels, suggesting SISU-102 is effective at inhibiting several viral kinetic gene classes. Indeed, since 72 h only allows for a single round of infection and SISU-102 was added at 24 hpi when IE proteins have reached maximum levels, these data suggest that SISU-102 is able to block lytic replication at a post IE stage in addition to limiting IE protein abundance. Accordingly, therapeutic inhibition of HSF1 significantly attenuated viral genome replication and progeny production similarly to ganciclovir treatment (**Fig. 5E, 5F**). Thus, the therapeutic inhibition HSF1 has the potential to accelerate the resolution of an active HCMV infection. Moreover, the ability of SISU-102 to attenuate HCMV replication post IE expression indicates HSF1 regulates multiple steps along the HCMV lytic replication cycle.

**Figure 5.**
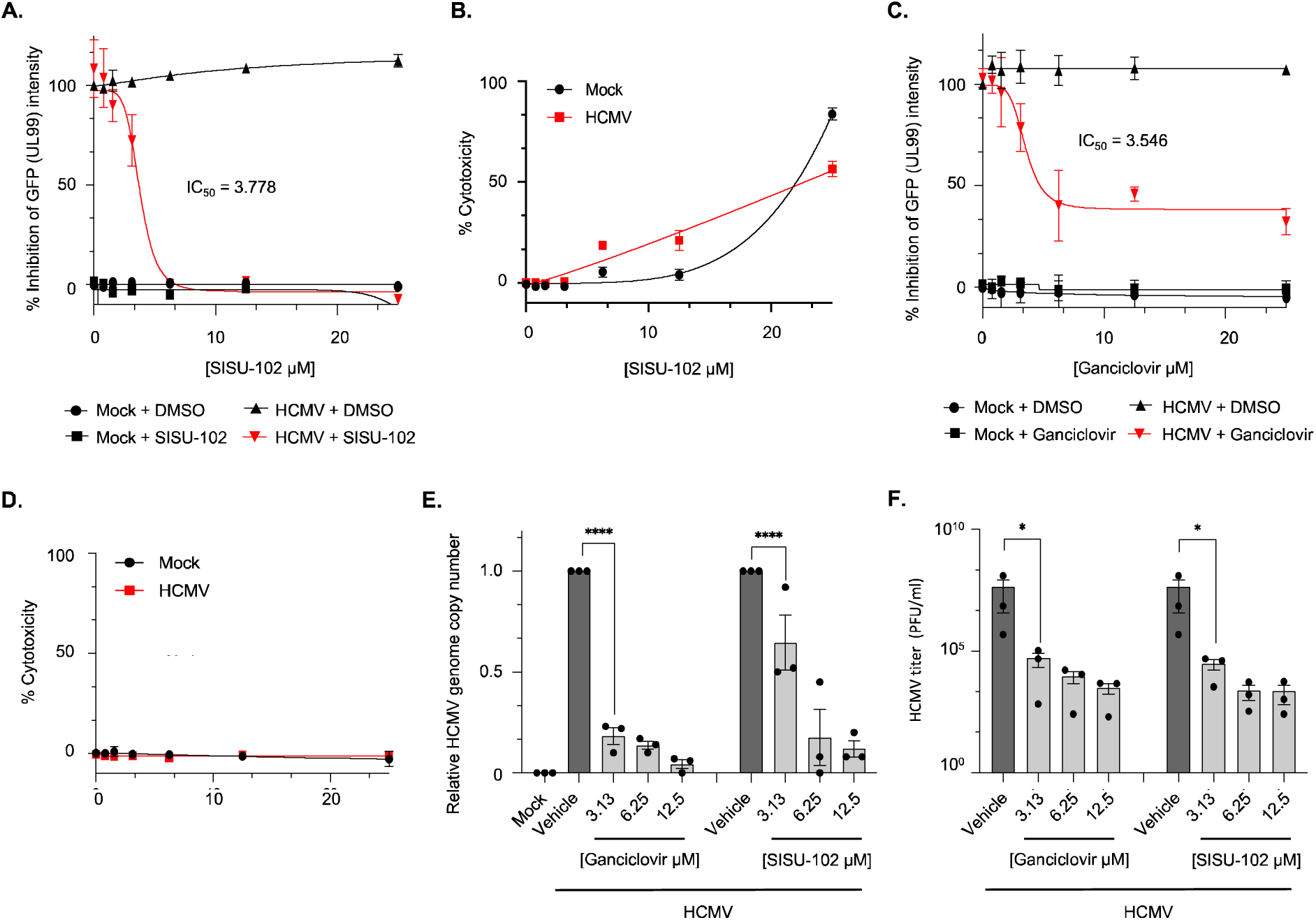
Therapeutic inhibition of HSF1 reduces HCMV L protein abundance and progeny production. **(A to F)** HEL 299 fibroblasts were infected (MOI 1) with UL99-GFP (A to D) or WT HCMV (E, F) for 24 h. Infected cells then prophylactically treated with SISU-102 or ganciclovir at the indicated concentrations for an addition 72 h. GFP intensity was measured using a fluorescent plate reader (A, C). Cytotoxicity was determined by SRB colorimetric assay (B, D). Viral genome copy number was assessed by qPCR analysis for *UL123* and *GAPDH* (E). TCID_50_ assays were performed on the supernatants to measure progeny virus production (F). All data are representative of 3 to 4 biological replicates per group. ****P < 0.0001, by one-way ANOVA with Tukey’s HSD post hoc test (E). *P<0.05, by Mann Whitney t-test (One-tailed) (F).

### Prophylactic inhibition of HSF1 attenuates HCMV replication and dissemination in a human skin xenograft transplant murine model

Due to the strict species specificity of HCMV, a human skin xenograft transplant murine model developed by our group to investigate in vivo HCMV lytic replication in biologically appropriate tissue under immunosuppressed conditions was utilized for the first time to assess the antiviral activity of SISU-102 during an acute infection (*42*). Human skin organ tissue provides a relevant setting for HCMV infection and contains many of the cell types critical for HCMV infection, including fibroblast, epithelial, endothelial, and myeloid cells (*42*). Human skin tissue from reduction mammoplasty was thinned, leaving the epidermis, dermis, and portion of the subdermal layer, which were implanted in nude athymic mice at 2 distal locations (**Fig. 6A**). After one month to allow for proper implantation and vascularization of the tissue, mice were prophylactically treated for 2 days with either drug carrier, SISU-102, or valganciclovir, a prodrug rapidly converted to ganciclovir (*43*). Following drug treatment, a single skin xenograft in each mouse was inoculated with PBS or a recombinant HCMV strain TB40E expressing GFP driven from a constitutively active SV40 promoter and a luciferase reporter gene replacing the nonessential UL18 glycoprotein driven from a true L promoter (ΔUL18-fLuc) (*42*). Measuring luciferase as a surrogate of L gene expression allows for continuous monitoring of HCMV replication by bioluminescence imaging of implanted skin xenografts. Because infection of tissue from different donors peaked at slightly shifted time points (12 to 14 days post infection), the day of peak infection was set as day 0 (D0) for all biological repeats to combine multiple data sets and perform subsequent statistical analyses. Initial infection of inoculated skin xenografts resulted in an early rise in bioluminescence in all treatment groups from -D13 to -D11 (**Fig. 6B**), suggesting the early increase in luciferase activity is not directly due to viral replication but likely from the delivery of luciferase packaged into the virion during viral egress. We found HCMV replication began at -D1, peaking at D0, and resolving by D2. SISU-102-treated mice infected with HCMV exhibited a significant reduction in viral replication at D0 beyond that of valganciclovir-treated mice, which is consistent with the increased in vitro potency of SISU-102 compared to ganciclovir at concentrations beyond their IC_50_’s (**Fig. 3A, 3C, 5A, 5C**). As expected, SISU-102 and valganciclovir were well-tolerated and had no adverse effect on body weight during the course of the study (**Fig. 6C**). Immunohistochemistry (IHC) analysis performed on skin xenografts at D2 confirmed increased GFP levels in the dermis area of inoculated xenografts, which was attenuated in mice prophylactically treated with SISU-102 or valganciclovir (**Fig. 6D**). To validate that the decrease in bioluminescence and GFP corresponded to decreased viral replication, D2 skin xenografts were treated with collagenase to get single cell dissociation and subjected to qPCR to measure viral genome copies or Western blotting to assess viral protein abundance. Both SISU-102 and valganciclovir significantly reduced viral genome copy numbers (**Fig. 6E**). Consistent with a decrease in viral replication, SISU-102 treatment reduced GFP as well as viral early (UL44) and late (gL) protein abundance (**Fig. 6F**). We also found that SISU-102 attenuated HSF1 levels within xenografts directly inoculated with HCMV, indicating HSF1 degradation activity by SISU-102 within infected xenografts. As expected, valganciclovir ablated gL abundance with minimal to modest effects on GFP and UL44 protein levels. Next, we examined the effects of SISU-102 on viral replication within distal uninoculated skin xenografts infected with virus disseminated from directly inoculated xenografts. We did not observe an early rise in bioluminescence at -D11 (3-5 dpi) in the uninoculated xenograft as with the directly inoculated site of infection, furthering supporting that the early increase in bioluminescence within the inoculated skin xenograft was from luciferase packaged within the virion. However, HCMV replication peaked at the distal uninoculated skin xenograft within the same timeframe as the inoculated xenograft at D0 (12-14 dpi) (**Fig. 6G**). The rapid dissemination of infected cells from the initial site of infection is consistent with infected myeloid cells migrating out of the inoculated tissue prior to 3 dpi (*42*). It should also be noted that circulating cell-free virus is unable to infect skin xenografts (*42*). Consistent with the suppression of viral replication within xenografts directly inoculated with HCMV, SISU-102 and valganciclovir prevented replication of disseminated virus within uninoculated skin xenografts. Thus, targeting HSF1 is a highly effective intervention at preventing in vivo viral replication and spread in an immunosuppressed setting. These findings further provide a proof-of-concept for the use of an in vivo human organ transplant murine model to test antivirals directed towards HCMV lytic replication in biologically relevant tissue.

**Figure 6:**
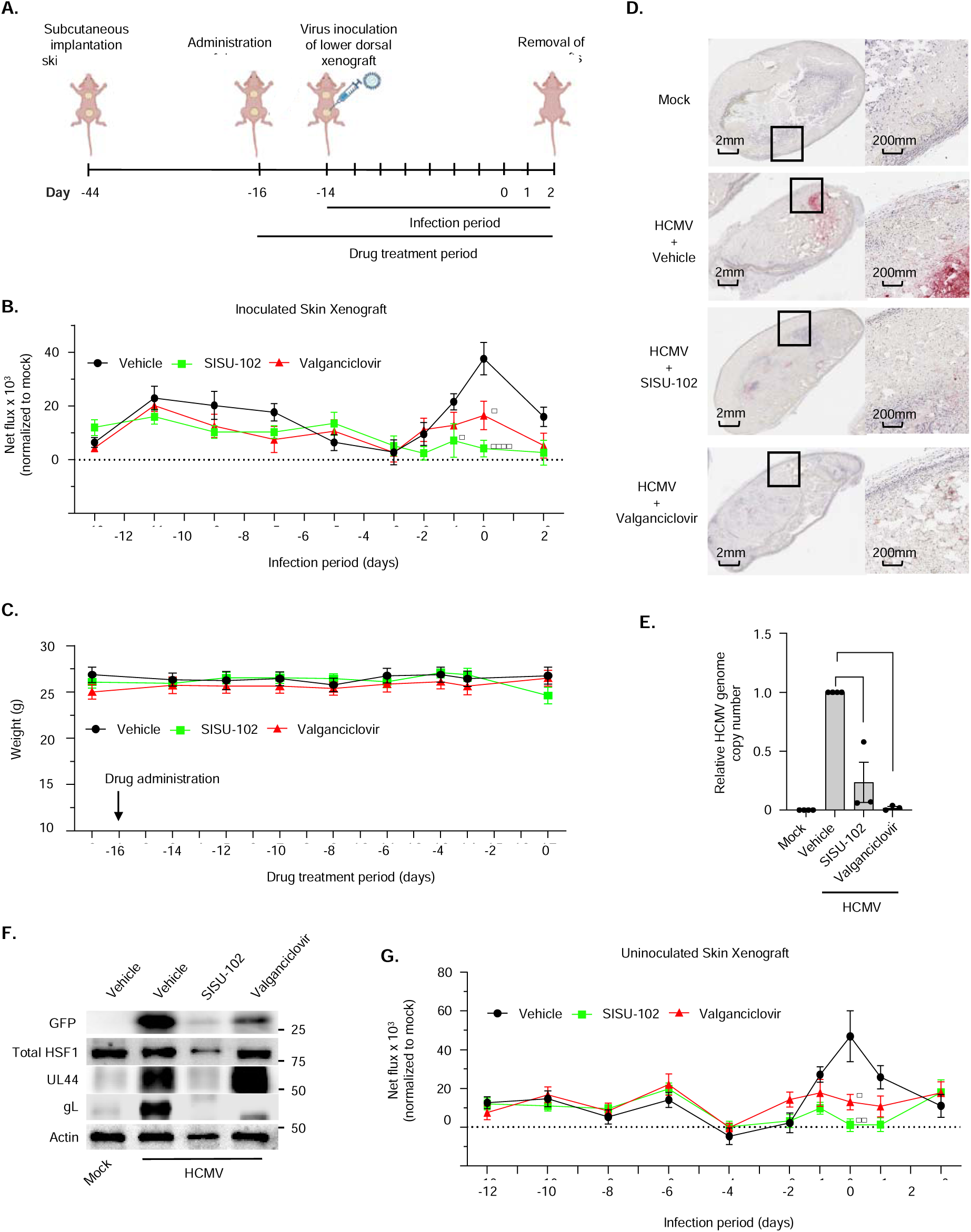
Prophylactic inhibition of HSF1 attenuates HCMV infection in vivo. **(A)** Timeline for skin implantation, drug treatment phase, and infection period of athymic mice. **(B to G)** Mice were treated daily beginning 2 days prior to infection with vehicle controls, SISU-102 (5 mg/kg/day), or valganciclovir (50 mg/kg/day). At day 2 following first drug administration, the bottom xenograft in each mouse was mock infected or infected with 10^6^ plaque forming units (pfu) of ΔUL18-fLuc. Bioluminescence imaging and quantification (total flux) of the inoculated (B) and uninoculated (G) skin xenografts were measured at the indicated time points using an IVIS Spectrum bioluminescence imaging system. Body weights of the mice were measured at the indicated time points throughout the drug treatment and infection phase (C). Bioluminescence and body weight data are representative of 7 to 22 mice per treatment group. Skin xenografts at day 16 post infection were collected, fixed, and processed for immunohistochemistry (IHC) to detect GFP (red) and nuclei (purple) (D). IHC data are representative of 5 biological replicates per group. At 16 days post infection, skin xenografts were harvested and homogenized into a single cell suspension (E, F). Viral genome copy number from each skin xenograft was determined by qPCR analysis for *UL123* and *GAPDH* (E). qPCR data are representative of 3 to 4 biological replicates per group. GFP, total HSF1, UL44, and gL were detected by Western blot (F). β-actin was used as a loading control. Western blots are representative of 5 biological replicates per group. *P<0.05, ****P < 0.0001, by one-way ANOVA with Tukey’s HSD post hoc test.

## DISCUSSION

Current FDA-approved antiviral drugs against HCMV target viral factors that mediate important steps along the lytic replication cycle. However, these therapeutics have several limitations, including adverse side effects and the emergence of drug-resistant strains. Ideally, novel antivirals targeting cellular factors that are specifically activated within infected cells and necessary for viral replication could alleviate the shortcomings of available anti-HCMV therapies. HSF1 is a stress response transcription factor hijacked by several viruses to promote the expression of viral genes (*25–29*), but whether HSF1 has any role during HCMV infection was unknown. Here, we show that HCMV entry into fibroblasts permissive for lytic replication stimulates a rapid activation of HSF1, as demonstrated by HSF1 nuclear accumulation and Ser^326^ phosphorylation, prior to viral gene expression. Using genetic knockdown approaches and SISU-102, which selectively targets and degrades nuclear HSF1 (*30*), we demonstrate that nuclear HSF1 is required for the expression of IE proteins. Importantly, SISU-102-mediated reduction of IE proteins led to a robust decrease in genome replication and virus progeny production similar to ganciclovir. To assess the in vivo efficacy of SISU-102, we leveraged a human skin xenograft transplant murine model developed by our group to examine acute HCMV infection in biologically relevant tissue under immunosuppressed conditions (*42*). Using this model as an antiviral testing platform for the first time, we showed the administration of SISU-102 suppressed in vivo replication in human tissue directly infected with HCMV, as well as prevented spread and replication in distal uninfected tissue with greater efficacy and potency than valganciclovir. Our study here demonstrates the essential role of HSF1 in promoting HCMV lytic replication and supports the targeting HSF1 as a host-directed antiviral therapy against HCMV infection.

HSF1 is an important cellular response against stress, yet our data indicate that the activation of HSF1 is not simply a cellular response to virus, but a well-orchestrated event controlled by HCMV. HS triggered a rapid and transient phosphorylation of HSF1, whereas HCMV stimulated a rapid and biphasic, chronic activation of HSF1 (**Fig. 1A to 1D**). Further, HSF1 activated during HS exhibited an increased molecular weight relative to HSF1 activated by HCMV infection at the same 30-min time point, suggesting distinct post-translational modifications induced during infection. These data demonstrate that HCMV stimulates a unique temporal activation of HSF1 with distinct biological activities. How HCMV modulates HSF1 activity remains unclear. UV-inactivated HCMV induced a rapid phosphorylation of HSF1 (**Fig. 1A, 1B**), indicating that viral entry alone is sufficient for the initial activation of HSF1. HCMV entry involves the binding of several viral glycoprotein complexes to cellular receptors to activate a multitude of signaling pathways necessary for the entry process. Many of these pathways are also known to regulate HSF1 phosphorylation, including the Akt, MAPK, mTOR, and AMPK protein kinase pathways (*44–46*). The induction of these pathways during viral entry is likely an evolutionary mechanism to rapidly activate HSF1 and “kick start” the transcription of IE genes. Once viral genes are transcribed, our data suggest that de novo synthesis of viral proteins is required to maintain HSF1 activity since infection with UV-HCMV infection induced only a transient activation (**Fig. 1A, 1B**). Whether the secondary activation is directly regulated by a viral gene product or is simply due to the accumulation of misfolded viral proteins within infected cells remains an active avenue of research. Regardless, our study shows that HCMV actively regulates HSF1 in virus-specific manner to promote the expression of viral genes that are required for virus production.

HSF1 mediates the transcription and expression of HSPs, many of which comprise cellular protein chaperones and quality control machinery to facilitate viral protein folding and assembly, and pro-survival proteins to maintain to integrity and function of infected cells, all of which serve to promote HCMV replication (*21–23*). Thus, HSF1 regulation of HCMV gene expression and genome replication could be mediated indirectly through HSPs and other cellular proteins. For instance, activated HSF1 enhances DENV replication by upregulating Atg7 (*27*). However, our data indicates an additional regulatory step. IE gene expression begins as early as 2 hpi, suggesting that the increased synthesis of HSPs may not occur in the timeframe needed to facilitate transcription of IE genes. Another possibility includes direct binding of HSF1 to viral promoters to initiate transcription of HCMV genes. The HSE consensus sequence consists of repetitive units of the nGAAn or nTTCn motif (*47, 48*). While the core nGAAn motif is relatively conserved, HSF1 binding can accommodate variations in the flanking regions making the sequence somewhat flexible and allowing HSF1 to regulate a wide array of genes (*48–50*). HSF1 directly binds to HSEs located in the viral promoters of HIV and EBV genes to promote the synthesis of viral proteins (*25, 28*). The presence of functional canonical or non-conical HSEs within the viral promoters of HCMV remains to be determined. Our data showing HSF1 regulates IE genes, which are transcribed within 2 hpi, strongly hints at HSF1 as directly regulating the promoter activity of HCMV genes. Thus, the targeting of HSF1 represents a potential host-targeted antiviral strategy to stall HCMV replication at multiple stages of infection.

The strict species specificity of HCMV makes in vivo testing of newly developed antivirals challenging. Current murine models rely heavily on fetal tissues to generate humanized mice (*51*) or the expression of human cells in a non-physiologically relevant environment, such as human fibroblasts placed in a three-dimensional matrix embedded into mice (*52*). Skin is a common site of HCMV reactivation in transplant recipients (*53*). Thus, we previously developed a human skin xenograft transplant murine model to allow for HCMV lytic replication in a physiologically relevant tissue under immunosuppressed conditions (*42*). Another advantage to this model is the ability to examine dissemination of HCMV infection away from the initial site of infection by the inclusion of a second transplanted skin xenograft at a distal site, which is infected by HCMV-containing myeloid cells originating from directly infected skin. Using this model, bioluminescence imaging showed both inoculated and uninoculated skin had similar infection kinetics (**Fig. 6B, 6G**), which is consistent with rapid migration and dissemination of infected myeloid cells out of infected tissue (*42*). Prophylactic SISU-102 treatment had slightly improved efficacy at suppressing HCMV replication at the initial inoculated tissue and at the disseminated site of infection when compared to valganciclovir, at much lower levels of compound. Previous studies also showed SISU-102 and HSF1 knockout to be well tolerated with no adverse effects on mice for weeks (*30, 54, 55*). Similarly, we found SISU-102 had no effect on weight, nor any observable side effects in both uninfected and infected mice over a 3-week period (**Fig. 6C**). These in vivo results, combined with the non-essentiality of HSF1 demonstrated in multiple mouse strains (*30, 54, 55*), suggest that the targeting of HSF1 selectively activated in HCMV-infected cells has the potential to be an effective and safe therapeutic intervention able to mitigate the emergence of drug resistance strains.

Overall, our study provides an important proof-of-concept for the use of HSF1 inhibitors as a novel antiviral therapeutic strategy to suppress HCMV replication. Because HSF1 regulates IE gene expression, which contrasts with current antivirals that only regulate post IE stages of lytic replication, these data also hint at the possibility of additive or synergistic effects with the combinational use of HSF1 inhibitors with current standard-of-care drugs. Further, there is the broader potential impact of the development of HSF1 inhibitors as antivirals since several viruses have evolved to utilize HSF1 to promote viral gene expression and viral replication, including HIV, dengue, EBV, and poxvirus (*25–29*). The ability to target multiple infections at once will greatly improve the prognosis of patients with co-infections, which commonly occur with opportunistic pathogens such as HCMV.

## MATERIALS AND METHODS

### Cell Culture

Human embryonic lung (HEL) 299 (CCL-137, ATCC) and MRC5 (CCL-171, ATCC) fibroblasts were maintained in Dulbecco’s Modified Eagle medium (DMEM) (Lonza) supplemented with 2.5 μg/ml plasmocin (Invivogen), and 10% fetal bovine serum (FBS) (MilliporeSigma).

### Virus Preparation and Infection

Construction of bacterial artificial chromosome (BAC)-derived TB40E-mCherry (wild-type; WT) (*56, 57*), TB40E-mCherry-IE2-*T2A*-eGFP (IE2-eGFP) (*38*), TB40E-mCherry-UL99-eGFP (UL99-eGFP) (*41*), and TB40E-eGFP-ΔUL18-fLuc (ΔUL18-fLuc) have been previously published or described below. All BACs were generously provided by Dr. Eain Murphy (SUNY Upstate Medical University) and Christine O’Connor (Cleveland Clinic). Viral BAC DNA and a pp71-expressing plasmid pCGN1-pp71 (1 μg) were transfected into MRC5 cells (CCL-171, ATCC). Cells were cultured in DMEM (Lonza) with 2.5 μg/ml plasmocin (Invivogen) and 10% FBS (MilliporeSigma). When 100% cytopathic effect was observed, cells were collected, lysed by sonication, and centrifuged (600 x *g*, 10 minutes (min), 22°C) to remove cellular debris. All passage 0 (P0) virus stocks were propagated on human embryonic lung (HEL) 299 fibroblasts (CCL-137, ATCC) in DMEM (Lonza) with 2.5 μg/ml plasmocin (Invivogen) and 10% FBS (MilliporeSigma). At 100% cytopathic effect, virus was purified from the supernatant by ultracentrifugation (115000 x *g*, 65 minutes (min), 22°C) through a 20% sorbitol cushion to remove cellular contaminants and resuspended in RPMI 1640 medium (ATCC). P1 virus was titered using 50% tissue culture infectious dose (TCID_50_) assays on HEL 299 fibroblasts (*58*) and used for all experiments. Levels of eGFP in fibroblast cultures infected with recombinant HCMV reporter viruses were measured using a BioTek Cytation 5 (BioTek). High passage HCMV strain Towne was also propagated as described above.

### Construction of **Δ**UL18-fLuc

Using a TB40E BAC engineered to express eGFP driven from a constitutively active SV40 promoter (TB40-eGFP) (*57, 59*), we constructed ΔUL18-fLuc by *galK* recombineering techniques, described in detail elsewhere (*60, 61*). Briefly, the *galK* gene was amplified from the pGalK plasmid by PCR using the primers listed in Table S1. Recombination competent SW105 *Escherichia coli* containing the TB40E-eGFP BAC were transformed with the resulting PCR product and *galK*-positive clones were selected. In parallel, firefly luciferase was amplified by PCR from pGL3-enhancer using primers listed in Table S1. Recombination competent SW105-containing *galK*-positive clones were then transformed with the resulting PCR product and counter selected against *galK*. The resulting virus, ΔUL18-fLuc, was validated by sequencing.

### Western Blot Analysis

Fibroblasts were harvested in modified radioimmunoprecipitation assay (RIPA) buffer (50 mM Tris-HCl pH 7.5, 5 mM EDTA, 100 mM NaCl, 1% Triton X-100, 0.1% SDS, 10% glycerol) supplemented with protease inhibitor cocktail (MilliporeSigma) and phosphatase inhibitor cocktails 2 and 3 (MilliporeSigma) for 30 min on ice. The lysates were cleared from the cell debris by centrifugation at 4°C (5 min, 21,000 x *g*) and stored at −20°C until further analysis. Protein samples were solubilized in Laemmli SDS sample nonreducing (6x) buffer (Boston Bioproducts) supplemented with β-mercaptoethanol (Amresco) by incubation at 95°C for 10 min. Equal amounts of total protein from each sample were loaded in each well, separated by SDS-polyacrylamide gel electrophoresis, and transferred to polyvinylidene difluoride membranes (BioRad). Blots were blocked in 5% bovine serum albumin (BSA) (Thermo Fisher Scientific) for 1 h at room temperature and then incubated with primary antibodies overnight at 4°C. The blots were incubated with horseradish peroxidase (HRP)-conjugated secondary antibodies (Cell Signaling Technology) for 30 min at room temperature, and chemiluminescence was detected using the Clarity Western ECL substrate (BioRad). Densitometry analysis was performed using Image Lab software (BioRad). The antibodies used in this study are described in Table S2.

### Subcellular Fractionation

Lysis gradients were made by layering of a mild lysis buffer onto a discontinuous iodixanol-based density gradient using OptiPrep density gradient medium (60% w/v iodixanol) (SigmaAldrich) for nuclear and cytoplasmic extraction as previously described (*62*). Briefly, live fibroblasts (2 x 10^6^ cells/sample) were loaded on top of the gradient followed by centrifugation (1000 x *g*, 10 min, 4°C). Cells travel through an initial wash buffer (8.34% OptiPrep) before entering a mild cell lysis layer (16.7% OptiPrep, 1% IGEPAL CA-630 (MilliporeSigma), 4 μg/ml Coomassie Brilliant Blue R250 (Biorad), 5 μg/ml Crystal Violet (MilliporeSigma)), which captures the cytoplasmic fraction. Undamaged nuclei pass through a subsequent nucleus wash layer (42% OptiPrep) prior to capture by a hyper-dense float layer (58.5% OptiPrep). Cytoplasmic and nuclear fractions were collected and prepared for Western blot analysis as described above.

### Sulforhodamine B (SRB) Cytotoxicity Assay

Cell viability was measured using the SRB Cell Cytotoxicity Assay (Abcam) as per the manufacturer’s instructions. Briefly, fibroblasts were seeded in 96-well plates at a density of 4 x 10 cells per well and cultured to confluency. Following treatment as described in the figure legends, cells were fixed with a fixation solution for 1 h at 4°C, washed 3x with ddH_2_O, and stained with the SRB protein-dye solution for 15 min protected from light. SRB dye was then removed, and the cells were washed 4x with a washing solution. Bound SRB was solubilized using the solubilization solution, and absorbance measured at 565 nm using a BioTek Epoch microplate reader (BioTek).

### siRNA Transfection

Fibroblasts were seeded at a density of 2 x 10 cells per well in 24-well plates and maintained in DMEM with 4% FBS in the absence of antibiotics. Cells were washed with 1x with PBS and transiently transfected with siRNAs using Lipofectamine RNAi/MAX reagent (Thermo Fisher Scientific) as per manufacturer’s recommendations. The following siRNAs were used: 100 nM of an HSF1-specific siRNA (Ambion; Catalog# AM16708) and 100 nM of a scrambled control siRNA (Invitrogen *Silencer*™ Negative Control No. 5; Catalog# AM4642). After siRNA transfection, fibroblasts were incubated for 48 h. Following the incubation period, cells were infected with HCMV for either 24 or 96 h. For the 96-h infection period, media was replaced after 24 h with fresh DMEM containing the corresponding siRNA to maintain gene silencing throughout the infection period.

### Human Skin Preparation

Adult female human breast skin was obtained from reduction mammoplasty surgeries in accordance with approved and established Institutional Review Board protocols and procedures at SUNY Upstate Medical University. All individuals donating tissue were healthy adults over 18 years old and gave consent per approved protocol (SUNY Upstate, Institutional Review Board). Skin tissue was collected in saline within 2 h of surgery and immediately processed (*42*). Whole-thickness skin tissue was cleaned with povidone-iodine and 70% ethanol to remove excess debris and washed with culture media. Skin was stretched tight across a Teflon-covered foam cylinder, secured with pins, and thinned using a Weck with a blade and Goulian guard to approximately 700 μm in thickness. The process of thinning the skin removes much of the underlying adipose and subdermal layer, leaving the epidermis and dermis intact. Thinned skin was cut into approximately 1-cm^2^ pieces to implant into mice.

### In Vivo Drug Testing

Thinned skin was introduced subcutaneously into 5-to 6-week-old athymic [CR ATH Ho; Crl: NU(NCr)-*Foxn1^nu^*] mice (Charles River) at 2 distal locations on the back near the head and tail of each mouse. A subcutaneous pocket was created by gently separating the skin from the underlying tissue using blunt dissection and a piece of skin inserted into the pocket with the epidermal side facing outward. Following implantation, skin xenografts were allowed to vascularize for 4 weeks. Mice were then treated daily during the drug treatment phase with either vehicle controls, SISU-102 (5 mg/kg/day by intraperitoneal injection; vehicle: 30% Captisol + 45% saline + 20% PEG 400 + 5% DMSO) (*27*), or valganciclovir (50 mg/kg/day oral gavage; vehicle: 0.5% methylcellulose, 0.5% Tween 80). 2 days after the drugs were first administered, the bottom xenografts were mock infected with Iscove’s Modified Dulbecco Medium (IMDM) (Lonza) or infected with ΔUL18-fLuc by directly injecting 10^6^ plaque forming units (pfu). For bioluminescence imaging, mice were administered with D-luciferin 150 mg/kg D-luciferin (MilliporeSigma) by intraperitoneal injection. After 10 mins, the mice were anesthetized with isoflurane and placed in the imaging chamber of an IVIS Spectrum bioluminescence imaging system (PerkinElmer). Bioluminescent signals from the infected skin grafts were quantified using Living Image software (PerkinElmer) and expressed as total flux (photons/second). At the end of the drug treatment phase, mice were euthanized, and skin xenografts were taken out for further processing to directly measure of viral replication. Single-cell suspensions of the harvested skin xenografts were prepared by homogenization of tissue using a glass homogenizer, followed by treatment with collagenase (1mg/ml) at 37°C until the tissue was fully dissociated into a single-cell suspension. Equal numbers of cells were used for subsequent qPCR and Western blot analyses.

### Immunohistochemistry (IHC)

Skin samples were fixed in 4% paraformaldehyde and sent to HistoWiz Inc. (histowiz.com) for processing using their Standard Operating Procedure and fully automated workflow. Briefly, samples were processed, embedded in paraffin, and sectioned at 4 µm. Immunohistochemistry was performed on a Bond Rx autostainer (Leica Biosystems) with enzyme treatment (1:1000) using standard protocols. An anti-GFP primary antibody (Abcam) was used, and detection was carried out using the Leica Biosystems DAB rabbit secondary polymer kit (Leica Biosystems). Bond Polymer Refine Detection system (Leica Biosystems) was used according to the manufacturer’s instructions. Red chromogen was used for final visualization and hematoxylin counterstaining was included as part of the standard procedure. After staining, sections were dehydrated and film coverslipped using a TissueTek-Prisma and Coverslipper (Sakura). Whole-slide scanning (40x) was performed on an Aperio AT2 (Leica Biosystems).

### Quantitative-PCR Analysis

Total DNA was isolated using the QIAamp DNA mini kit (Qiagen) as per the manufacturer’s recommendation. For quantitative PCR, *UL123* and *GAPDH* were detected with CFX Connect Real-Time System (Bio-Rad) using iTaq Universal SYBR Green Supermix (Bio-Rad) as per the manufacturer’s recommendation. Samples were analyzed in technical duplicates and normalized to GAPDH. Oligonucleotide sequences used in this study are described in Table S1.

### Statistics Analysis

All experiments were performed with a minimum of 3 biological replicates per group. Data were analyzed using one-way ANOVA with Tukey’s honest significant difference (HSD) post hoc test with GraphPad Prism software, and *p*-values less than 0.05 were considered statistically significant.

## List of Supplementary Materials

Figs. S1 to S2

Table S1 and S2

## Acknowledgments

We thank Christine Burrer in the Department of Microbiology and Immunology at SUNY Upstate Medical University for technical support, maintenance of lab operations, and assistance with virus growth and isolation.

## Author contributions

Conceptualization: GCC, DJT, DA

Methodology: DA, JB, MJM, JFM, EAM, CMO

Investigation: GCC, DA

Visualization: GCC, DA

Funding acquisition: GCC

Project administration: GCC

Supervision: G.C.C.

Writing – original draft: GCC, DA

Writing – review & editing: GCC, DJT, DA

## Data and materials availability

All data needed to evaluate the conclusions in the manuscript are present in the paper or the Supplementary Materials. All compounds are commercially available. Recombinant viruses are available from G.C. under a material transfer agreement with SUNY Upstate Medical University, Syracuse, NY 13210, USA

