## Supplemental Material for "Targeting the host transcription factor HSF1 prevents human cytomegalovirus replication in vitro and in vivo"

Dilruba Akter et al.

**This PDF file includes:**

Figs. S1 and S2

Table S1 and S2

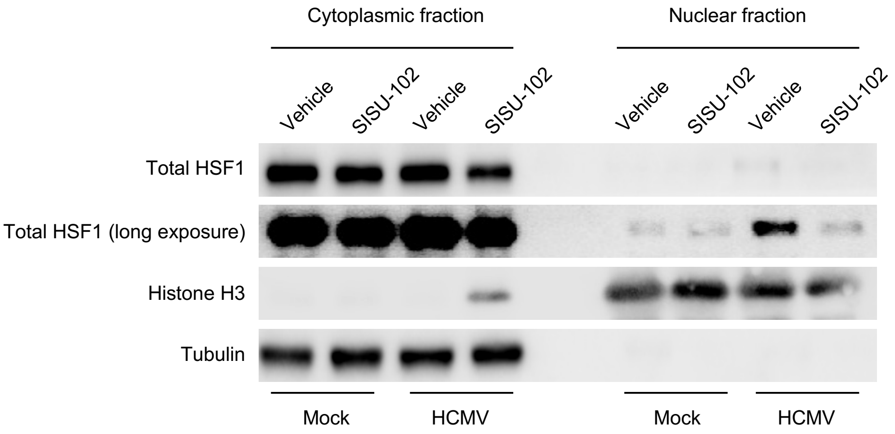


**Fig S1. HCMV infection drives HSF1 into the nucleus.** HEL 299 fibroblasts were treated with vehicle or 6.25 μM of SISU102 for 24 h. Cell were then mock infected or infected (MOI 5) with WT HCMV for 24 h. Live fibroblasts layered onto a discontinuous iodixanol-based gradient and centrifuged to separate soluble cytosolic proteins and nuclei. Cytoplasmic and nuclear HSF1 was detected by Western blot. Tubulin and histone H3 served as cytosolic and nuclear loading controls, respectively. Western blots are representative of at least 3 biological replicates per group.

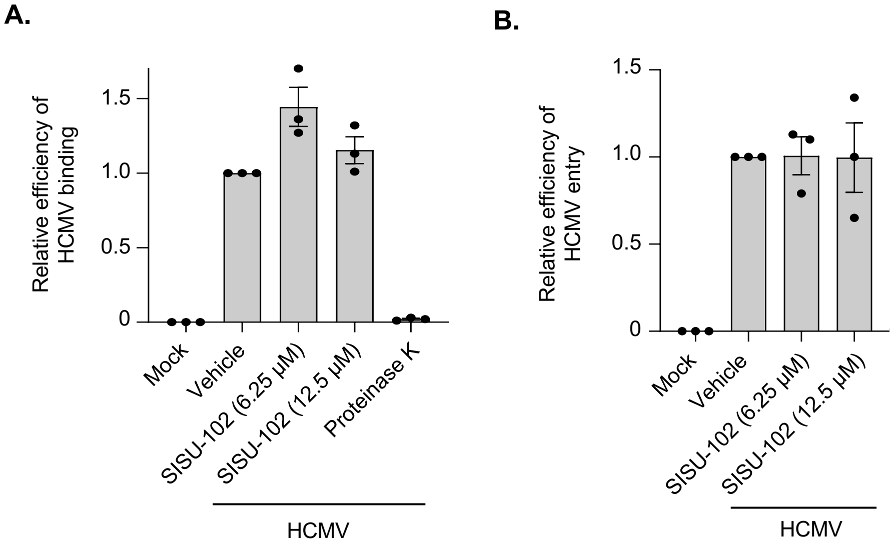


**Fig S2. Inhibition of HSF1 does not affect HCMV binding and entry into fibroblasts. (A, B)** HEL 299 fibroblasts were treated with vehicle or SISU102 at the indicated concentrations for 24 h. **(A)** Cells were then incubated on ice for 1 h and infected with WT HCMV for additional 2 h on ice to allow for viral binding. As a control to ensure HCMV remained at the cell surface, infected fibroblasts were treated with 1 mg/ml proteinase K (PK) for an additional hour on ice to strip off surface bound virus particles. Cells were washed with PBS to remove any unbound viral particles. **(B)** To assess viral entry, parallel samples were temperature shifted to 37$^{o}$C for 2 h following viral binding to allow for internalization of the bound virus particle. Cells were then treated with 1 mg/ml PK for 5 min at 37$^{o}$C and washed with PBS to remove surface bound virus particles not internalized into the cell. Viral binding and entry were measured by qPCR analysis for *UL123* (as a marker of the viral genome) and *GAPDH*. Data are means ± SEM from 3 biological replicates per group.

**Table S1. Oligonucleotides used in this study**

| Primer Use | Sequence¶ | Orientation |
|---|---|---|
| *galK* insertion | GAAAGTATATAACGCCGATCATGTCCGAGGAACTGTTAATAAAACGCCATGCCTGTTGACAATTAATCATCGGCA | forward |
| | CGGGGGCACGCGGTAACCGACGTCGAAACAGCTCATACAGGGCGTTGCTGGTCAGCACTGTCCTGCTCCTT | reverse |
| Luciferase (UL18) insertion | GAAAGTATATAACGCCGATCATGTCCGAGGAACTGTTAATAAAACGCCATG**ATGGAAGACGCCAAAAACAT** | forward |
| | CGGGGGCACGCGGTAACCGACGTCGAAACAGCTCATACAGGGCGTTGCTGG**TTACACGGCGATCTTTCCGC** | reverse |
| Sequencing primers | TGATGTCGGATCCGGCACGGT | forward |
| | CCAACACGAACATAGTGTAGT | reverse |
| *UL123* | AGTGACCGAGGATTGCAACG | forward |
| | CCTTGATTCTATGCCGCACC | reverse |
| *GAPDH* | ACCCACTCCTCCACCTTTGAC | forward |
| | CTGTTGCTGTAGCCAAATTCGT | reverse |

¶Primer sequences are shown 5’ to 3’. Underlined sequences denote those corresponding to the pGalK plasmid. Bolded sequences denote those corresponding to the pGL3-enhancer vector (Promega), from which firefly luciferase was amplified.

**Table S2. Antibodies used in this study.**

| Antibody | Dilution | Company (Catalog) |
|---|---|---|
| β-Actin (Rhodamine conjugated) | 1:2000 | BioRad (12004163) |
| p-HSF1 ($Ser^{326}$) | 1:500 | Abcam (ab76076) |
| GFP | 1:100 | Abcam (ab183734) |
| Total HSF1 | 1:1000 | Cell Signaling Technology (12972S) |
| α-tubulin | 1:1000 | Cell Signaling Technology (3873S) |
| Histone H3 | 1:2000 | Cell Signaling Technology (4499S) |
| UL44 | 1:1000 | MyBioSource (MBS530793) |
| IE1 | 1:1000 | A generous gift from Dr. Eain Murphy (SUNY Upstate Medical University) |
| gL | 1:500 | A generous gift from Dr. Eain Murphy (SUNY Upstate Medical University) |
